# A 3D cell-free bone model shows collagen mineralization is driven and controlled by the matrix

**DOI:** 10.1101/2022.10.24.513466

**Authors:** Robin H.M. van der Meijden, Deniz Daviran, Luco Rutten, X. Frank Walboomers, Elena Macías-Sánchez, Nico Sommerdijk, Anat Akiva

**Author notes:** Geert Grooteplein Zuid 28 Nijmegen, 6525 GA, The Netherlands.

## Abstract

Osteons, the main organizational components of human compact bone, are cylindrical structures composed of layers of mineralized collagen fibrils, called lamellae. These lamellae have different orientations, different degrees of organization and different degrees of mineralization where the intrafibrillar and extrafibrillar mineral is intergrown into one continuous network of oriented crystals.

While cellular activity is clearly the source of the organic matrix, recent in vitro studies call into question whether the cells are also involved in matrix mineralization, and suggest that this process could be simply driven by the interactions of the mineral with extracellular matrix.

Through the remineralization of demineralized bone matrix, we demonstrate the complete multiscale reconstruction of the 3D structure and composition of the osteon without cellular involvement. We then explore this cell-free in vitro system as a realistic, functional model for the in situ investigation of matrix-controlled mineralization processes. Combined Raman and electron microscopy indicates that glycosaminoglycans play a more prominent role than generally assumed in the matrix-mineral interactions. Our experiments also show that the organization of the collagen is in part a result of its interaction with the developing mineral.

## 1. Introduction

The main organizational components of compact adult human bone are cylindrical structures called osteons.^[1]^ The walls of these osteons, consist of organized layers of collagen fibrils named lamellae, that are mineralized with plate-like crystals of carbonated hydroxyapatite (cHAp). These mineralized collagen fibrils are organized over multiple hierarchical levels,^[2]^ where the subsequent layers have different orientations, different degrees of organization and different degrees of mineralization^[3]^ (Figure 1a). Moreover, the intrafibrillar and extrafibrillar mineral is intergrown into one continuous network of oriented crystals.^[4]^

**Figure 1.**
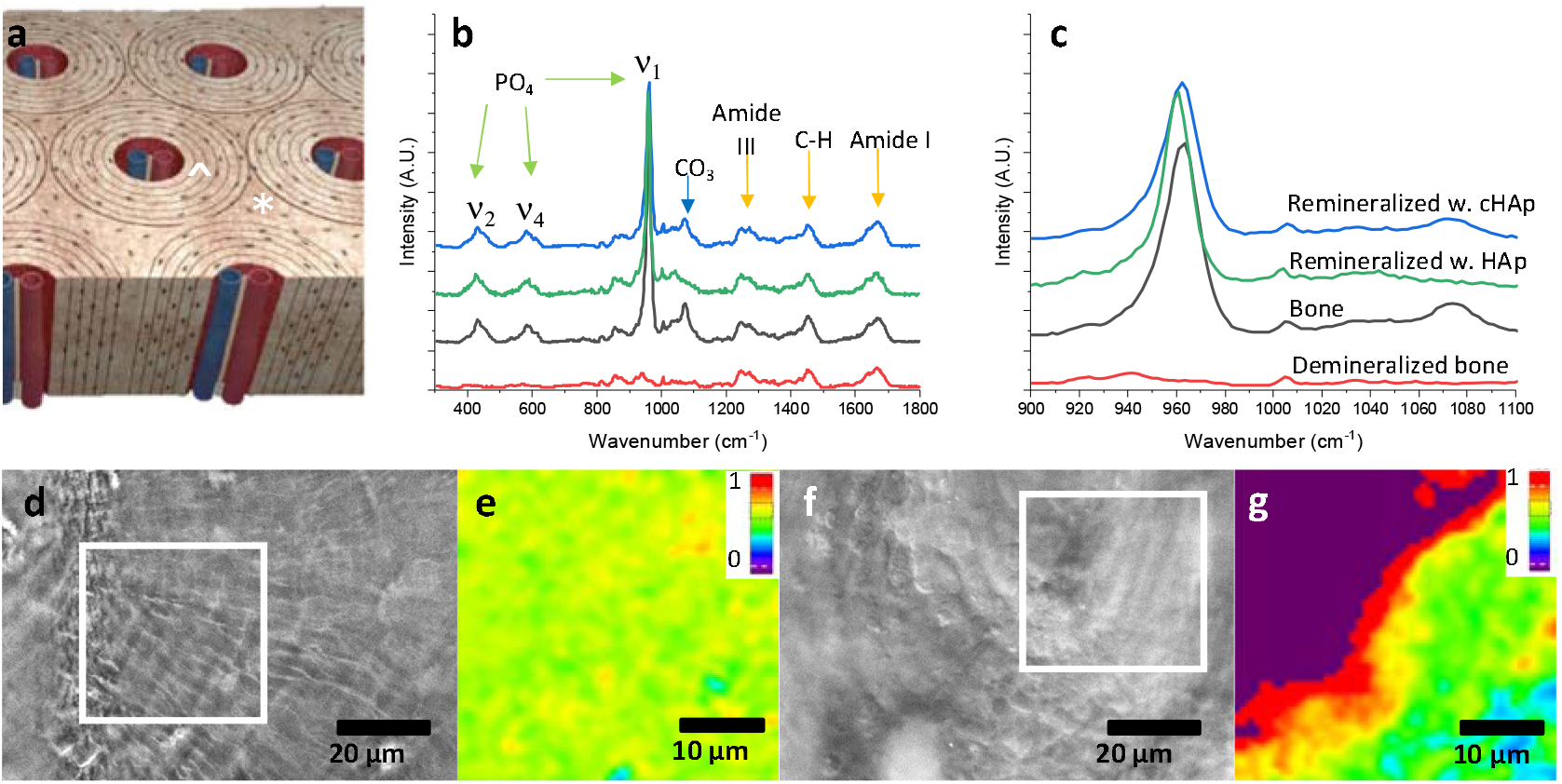
Remineralization of bone extracellular matrix. a) Schematic of the cortical bone, indicating the osteons (^) and interstitial bone (*). b) Raman spectra of demineralized bone (red), the corresponding control bone (black), the matrix re-mineralized with HAp (green), and matrix re-mineralized with cHAp (blue). c) Zoom in on the main mineral peak in (b) showing the change in the position and width of the v1 PO_4_ peak in HAp (green) related to carbonate substitution (blue). d, f) Grey scale optical images of (d) control bone and (f) the re-mineralized matrix, showing the lamellae of osteons. e, g) Raman color maps recorded in the area marked in white boxes in (d) and (f) indicating the mineral-to-matrix ratio distribution in (e) the control bone and (g) the re-mineralized matrix, based on the intensity ratios of the ν2 PO_4_ (431 cm^-1^) and CH_2_ vibrations (1450 cm^-1^). The large green area in (g) shows that the level of mineralization achieved in the bulk is in the range of natural bone as shown in (e). The purple area in g) indicates the location of the haversian canal.

The osteons are created and maintained through the interplay of three bone-specific cell types: osteoclasts (bone removing cells), osteoblasts (bone forming cells) and osteocytes that direct the activity of both.^[5]^ Osteoblast activity is clearly responsible for the production of the osteoid, i.e. the organic 3D organized collagenous matrix, but recent studies call into question whether the cells are also actively involved in controlling the mineralization of the collagen matrix.^[6]^ Although the current prevailing thought is that osteoblasts concentrate, package and transport the mineral to the calcifying collagen matrix in the form of calcium phosphate loaded vesicles,^[7]^ recent evidence shows that in developing bones, direct connections of the vasculature to the mineralization front could explain fast exchange of material.^[8]^ This points at the possibility that mineral is supplied directly from the blood, with no necessary involvement of the osteoblasts in the mineralization process during the early stages of bone development.^[8e, 9]^

Specifically, *in vitro* experiments have demonstrated that intrafibrillar mineralization of collagen can be achieved without cellular involvement in the presence of crystallization inhibiting macromolecules (e.g. poly(aspartic acid),^[10]^ fetuin^[10a]^ and osteopontin^[11]^) yielding mineralized fibrils that are indistinguishable from those in bone,^[6a, 12]^ and that confinement by the collagen is an essential factor in directing mineralization.^[6a]^ Beyond the mineralization of individual collagen fibrils, cell-free in vitro experiments^[13]^ demonstrated the remineralization of thin sections of demineralized periodontal tissue from mouse molars with control over the 2D organization of the collagen crystals within the collagen matrix,^[14]^ while 3D bulk samples of demineralized manatee bone showed the formation of aligned mineral over depths of 100 *µ*m.^[10b]^

Based on the above we hypothesize that bone biomineralization is driven solely by solution composition and matrix-mineral interactions. The organic bone matrix consists of the hierarchically organized collagen modified with carbohydrates and supplemented with non-collagenous proteins.^[15]^ In the last decades, much attention has been given to the role of non-collagenous proteins (NCPs) in bone biomineralization, and for many of them roles have been assigned.^[16]^ However, more recent studies showed that NCPs were not essential for collagen mineralization,^[17]^ and others pointed to the importance of matrix modifications with carbohydrates,^[18]^ and in particular glycosaminoglycans (GAGs),^[19]^ in local control over mineralization. Despite their importance, the role of the different GAGs in regulating bone biomineralization is still not well understood and for similar systems both inhibitory^[20]^ and promoting^[19a]^ effects have been reported.

In the present work we demonstrate the complete reconstruction of the 3D hierarchical multiscale mineral structure of the osteon and its native chemical composition without cellular involvement. We then explore this cell free *in vitro* system as a realistic, functional model for the *in situ* investigation of matrix controlled mineralization processes, focusing on the collagen-mineral interplay and the role of GAGs in controlling collagen mineralization.

We use a combination of electron microscopy and Raman microscopy to demonstrate the successful *in vitro* remineralization of human bone extracellular matrix (ECM), regaining the original chemical composition of native bone, while reconstructing the 3D hierarchical structure of the osteon from the level of intrafibrillar collagen mineralization, to the formation of the 3D intertwined mineral network and to the oriented lamellar structure. We also demonstrate that this cell-free *in vitro* system recapitulates several features of the bone mineralization process previously reported from *in vivo* and *ex vivo* studies. ^[4, 9, 19a, 21]^ Finally, using live *in situ* monitoring of these processes we show how collagen structure is imposed during mineralization, and propose that the presence of glycosaminoglycans may play a role in controlling the local rate of mineralization, but also the crystallization pathway by which the mineral is formed.^[21b, 22]^

## 2. Results

### Reconstructing the ECM chemistry and structure from demineralized tissue

#### Reconstructing the bone ECM Chemistry

Cortical bone samples from human femur shafts were divided into two parts, where one was demineralized and the other one used as a control. Complete demineralization was confirmed with Raman spectroscopy (Figure 1b,c), inductively coupled plasma optical emission spectroscopy (ICP-OES), X-ray microscopy, attenuated total reflection – infrared spectroscopy (ATR-FTIR) and scanning electron microscopy (SEM) (Figure S1). To demonstrate reconstitution of the original bone structure upon remineralization, pieces of the demineralized sample were submerged in an biomimetic mineralization solution (0.8x PBS containing 1.7 mM CaCl_2_, 1.0 mM KH_2_PO_4_) at 37°C using 100*µ*g/ml poly(aspartic acid) (pAsp) as crystallization control agent^[10a, 10c, 14]^. After 4 days, combined Raman spectroscopy (Figure 1b,c,) and Raman microscopy (Figure 1d-g) confirmed the infiltration of the matrix with hydroxyapatite (HAp). Comparison with the control showed that the mineral-to-matrix ratio in the investigated areas had again reached the natural levels, after 4 days over large areas of mineral infiltration, as judged from the ratios of the bands associated with the organic matrix (1450 cm^-1^, CH_2_-stretch) and the mineral (431 cm^-1^ PO_4_ ν2) (Figure 1e,g, and Figure S2, S3), These levels did not significantly increase further upon prolonged exposure to the mineralization solution, and after 21 days still mineralization levels similar to natural bone were obtained (Figure S3). Small changes were observed in the main mineral peak position (PO_4_, ν1; 959 cm^-1^) and width (FWHM 14 cm^-1^) compared to the control (961 cm^-1^ and FWHM 18 cm^-1^, respectively). We attribute these differences to the absence of carbonate substitution in the re-mineralized sample, which reaches 8-10% in natural bone tissue.^[23]^ Our assessment was confirmed by the formation of cHAp through the addition of 25 mM of HCO_3_^-^ to the mineralization solution (Figure 1b). Raman spectroscopy indeed indicated natural carbonate substitution levels as judged from the intensity of the carbonate ν2 vibration (1072 cm^-1^). Moreover, the main PO_4_ ν1 peak had indeed regained the original position (961cm^-1^) and width (FWHM 17 cm^-1^) (Figure 1c) resulting in a near perfect spectral match between the re-mineralized sample and the control. This implies that the composition of the bone mineral phase can be fully regenerated in this *in vitro* mineralization system.

Considering our aim to investigate the role of GAGs in the mineralization process, and their low abundance in the bone matrix,^[19b]^ it is important to note that there is significant spectral overlap between the main (but weak) peaks of the GAGs with the carbonate peak of the mineral (1072 cm^-1^), and both the amide I (1666 cm^-1^) and amide III (1250 cm^-1^) peaks of the collagen (Figure S4). Here, the carbonate-free remineralization of the matrix provided an opportunity: remineralization with a carbonate free mineralization solution would provide a more open spectral window between 1050 and 1800 cm^-1^ that would aid the identification of GAG related signals in the Raman spectra. It was therefore decided to re-mineralize the matrix with pure HAp, rather than cHAp.

#### Reconstructing the bone nano-structure

Although the above results demonstrate that the chemical composition of the bone extracellular matrix can be achieved by remineralization of the osteoid - without further involvement of cellular activity - they do not show whether these cell-free *in vitro* experiments can also recapture the multilevel hierarchical structure of the mineralized matrix. Recently the bone extracellular matrix was shown to consist of intimately interpenetrating collagenous and mineral phases, in which a cross-fibrillar network of intergrown intrafibrillar and interfibrillar mineral platelets provides structuring from the nanoscale to the multi-micron level.^[4]^

To certify that cell-free osteoid remineralization also with high fidelity reconstitutes this above nano-to-meso structural architecture of bone, thin sections (200 nm thick) of the demineralized bone were place on gold TEM grids and exposed to the mineralization solution. TEM analysis revealed that the collagen matrix indeed showed an organized meso-structure indistinguishable of that of bone, having lamella of collagen in different orientations with respect to the viewing direction (Figure 2a), similar to previous findings.^[4, 24]^ At higher magnification the mineralized collagen showed both the so-called lacey (Figure 2b) and the lamellar (Figure 2c) patterns that are typical for the bone ultrastructure when the mineralized collagen fibrils are oriented perpendicular and parallel to the viewing direction, respectively.^[4, 24]^ Selected area electron diffraction of the lamellar structure (Figure 2d) confirmed that the mineral phase consisted of apatite crystals that were aligned with their c-axis along the long axis of the collagen fibril. As in bone, the (002) and (004) reflections showed arcs with an angular dispersion of ± 25 degrees, showing an uniaxial orientation of the crystals. The unresolved reflections corresponding to (211), (112), (300), (202) and (301) appear combined in a diffraction ring.^[4, 6a, 25]^

**Figure 2.**
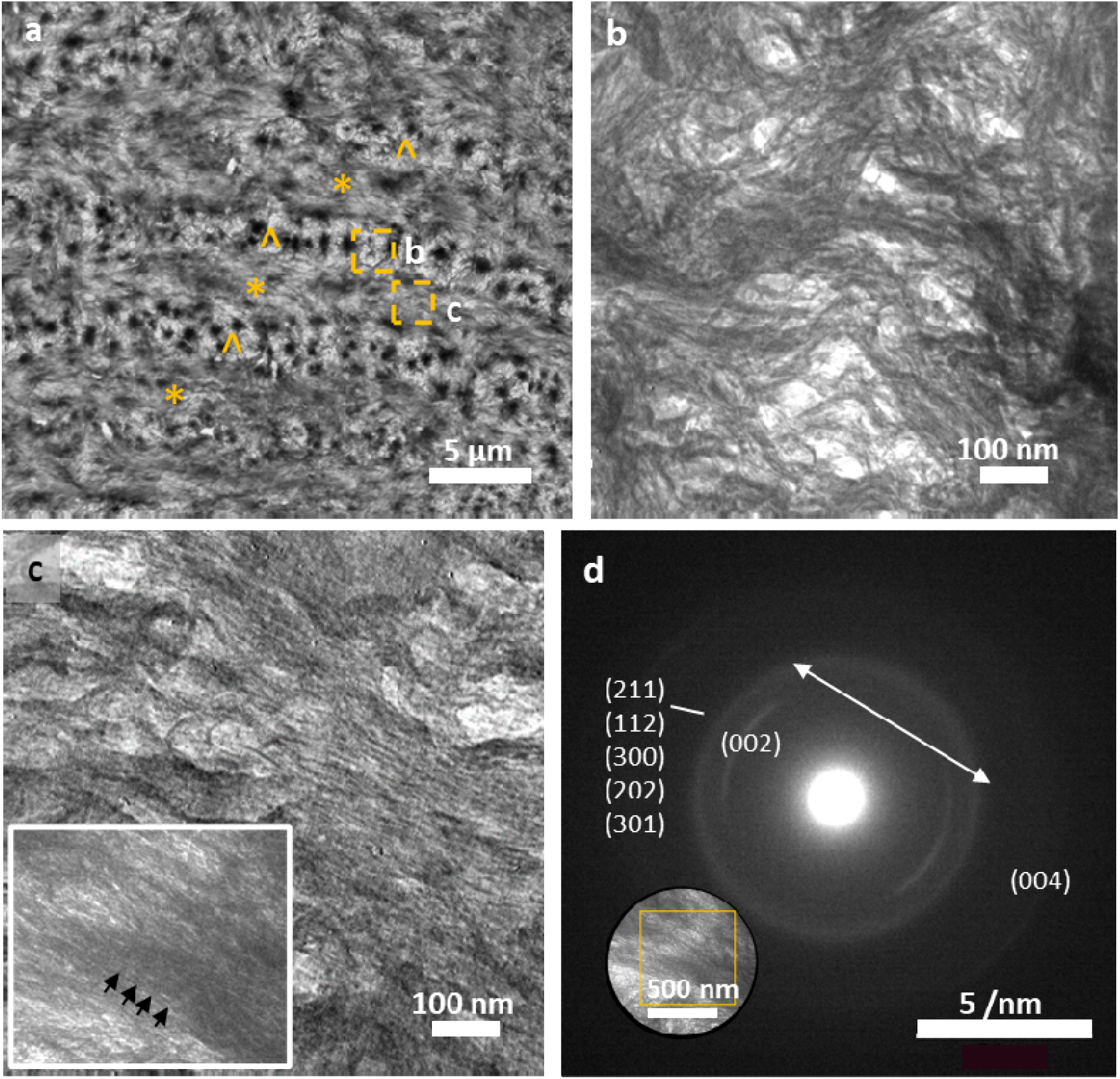
TEM of the re-mineralized matrix. a) a large overview showing differently oriented lamella of parallel(*) and perpendicularly(^) oriented collagen fibrils. Boxes indicated areas where high magnification images in b-c were taken. b) Lacey pattern representing the perpendicularly oriented collagen, similar as reported for natural bone ECM. c) Parallel collagen fibrils with mineral crystallites oriented along the long axis of the fibrils. Inset: overview with arrows highlighting the characteristic d-banding pattern of the collagen. d) Selected area electron diffraction pattern taken from the area in the inset showing a typical diffraction pattern for bone apatite. The arcs of the (002) reflections indicate the c-axes of the crystals are oriented along the collagen direction (arrows). Insets in (c) and (d) show the same area.

#### Reconstructing the hierarchical structure of bone

After establishing that this system restored both the chemical composition and the nanostructure of the bone, we set out to demonstrate that it also can reconstruct the next hierarchical level, i.e. the 3D structure of the osteons, in which concentric micron-sized layers of parallel oriented mineralized fibrils form either a so-called plywood-like structure^[21a]^ where each layer is rotated with respect to the neighboring one or a fanning array pattern in which the fibrils have a more gradual change in their orientation.^[26]^ To investigate the faithful regeneration of this sub-millimeter structure by *in vitro* mineralization of the osteoid, polarized Raman microscopy was used to probe the orientation of the mineral in the different concentric layers of collagen fibrils, using the original unmodified sections of the same bone tissue as a control.

As the Raman vibrational modes of both the phosphate ν1 (961 cm^-1^) and amide I (1666 cm^-1^) peaks in bone samples are orientation dependent, polarized Raman microscopy can independently visualize their orientation by mapping the peak intensities normalized to the polarization independent Amide III peak (Figure S6).^[26]^ We used this to map the alignment of the mineral with respect to the collagen matrix throughout the different layers of the osteon (Figure 3a,b).

**Figure 3.**
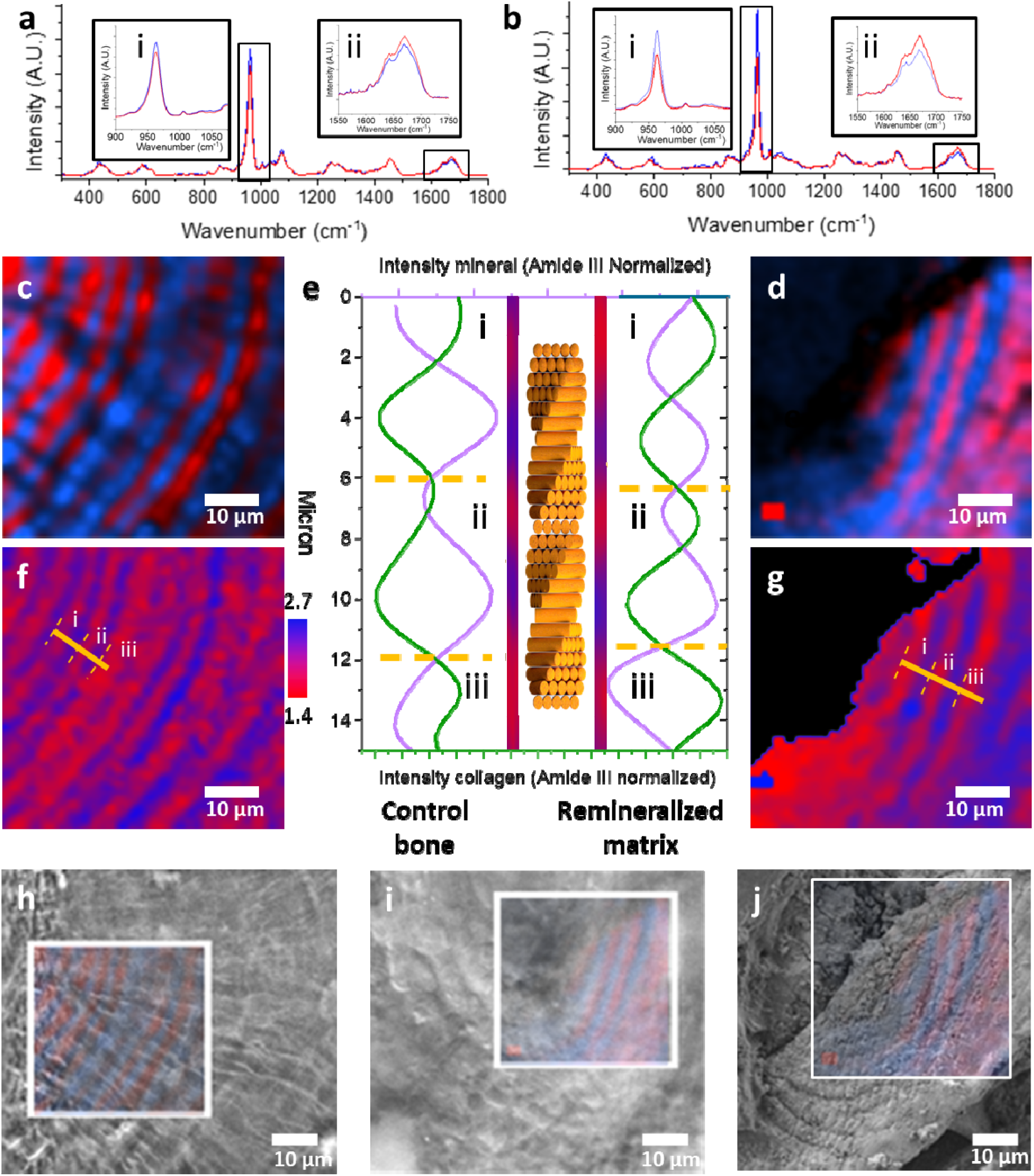
Polarized Raman microscopy shows restauration of the 3D osteon structure. a,b) Spectra and c,d) corresponding maps from component analysis of the Raman data in figures 1d,f). a,c) control bone and b,d) re-mineralized matrix. Color coding of the spectra matches the colors of the component maps. Insets and boxes in a,b) highlight the intensity variation of orientation dependent peaks of i) the ν_1_ PO_4_ and ii) the amide I vibrations while using a fixed laser orientation, demonstrating that the components in a,c) represent a difference in orientation of both the collagen and mineral over the different lamellae. e) Intensities of the individual ν_1_ PO_4_ (purple) and the amide I (green) peaks after normalization to the orientation independent Amide III vibration, representing independently the orientations of the mineral and organic phase over the orange line in f,g). For the mineral a maximum corresponds to a parallel alignment of the c-axis with the laser direction, whereas for the organic phase a maximum represents a perpendicular organization of the fibril to the laser direction. Central inset shows a schematic of the corresponding orientation of the collagen bundles. The red-blue stripes adjacent to the schematic are extracted of the orange lines in f,g, showing the correlation of the imaging with the spectral data. Dotted lines indicate the ranges of a full rotation of the collagen molecules, in total both lines show 2,5 periods. f,g) Map of the ratio of the ν_1_ PO_4_ and the amide I intensities, showing their gradual, inversely coupled changes in rotation over the different lamellae. h-j) Overlay of the Raman microscopy images with optical and scanning electron micrographs showing chemical and structural correlation of the lamellae. h) Overlay of Raman map shown in c) with optical micrograph in Fig 1d, i) Overlay of the Raman map shown in d) with optical micrograph in Figure 1f, j) overlay of the Raman map shown in d) with SEM image of the remineralized matrix.(see also S5)

Unsupervised component analysis (see S2) of the Raman maps identified, for both the re-mineralized sample and the control bone, two spectral components indicating different ratios of the peak areas of the phosphate ν1 and amide I (Figure 3c,d). These components represent two principal orientations (parallel and perpendicular to the imaging plane) of the mineralized collagen fibrils in the different lamellae of the osteon.^[27]^ However, analyzing the absolute intensities of the individual amide I and phosphate ν_1_ vibrations values of each pixel revealed for both samples a continuous sinusoidal variation through the different layers (Figure 3e), rather than an alternating perpendicular organization as is implied by the component analysis. Comparing the intensities of the two vibrations (phosphate ν_1_ and amide I) revealed an opposite angular dependence (i.e. where the mineral peak increases, the collagen peak decreases and *vice versa*), demonstrating that the orientation of the mineral is indeed coupled to the orientation of the collagen fibrils. The sinusoidal variation in orientation (pitch ∼6 *µ*m) furthermore demonstrates that in the investigated bone material the collagen fibrils are arranged in a fanning array structure with a layer thickness of < 500 nm (limited by the map resolution, Figure 3e,f,g).^[26]^ rather than in a twisted plywood structure which has sharp rotational offsets of ∼90 degrees between aligned collagen layers that also has been reported, as the latter would have sharp transitions between two orientations.^[21a]^

### A cell-free *in vitro* model for bone biomineralization

The above shows that re-mineralization of the collagenous matrix can faithfully restore both the chemical and the hierarchical structure of the bone extracellular matrix - from the nanoscopic intrafibrillar mineral orientation to the sub-millimeter concentric lamellae of alternatingly oriented mineralized collagen fibrils in the osteon. As matrix mineralization had not yet been completed throughout the entire sample, no bulk measurements analyzing mineral content or mechanical properties were performed. Nevertheless, the above results imply that our model follows the mineral infiltration and maturation pathways observed during bone biomineralization, and hence can be used to study the effect of local matrix modifications on this process.

To establish this, we provided the demineralized samples with a fixed amount of mineralization solution which would deplete during the reaction,^[28]^ thereby trapping intermediate stages of the process. Accordingly, demineralized bone samples were incubated for 8 days in a small amount of biomimetic mineralization solution, with only one renewal of the solution after 4 days. SEM with back scatter electron (BSE) detection indeed showed a partially re-mineralized bone matrix in which isolated high intensity (mineral) areas were observed (Figure 4a), similar to observations in developing mouse bone (Figure 4b, Figure S6). ^[8b, 8c, 9, 29]^ BSE intensity profiles revealed a sharp peak for the unmineralized collagen and a broad overlapping peak of the mineralizing collagen (Figure 4c), indicating a distribution of mineral density within the mineralizing matrix.

**Figure 4.**
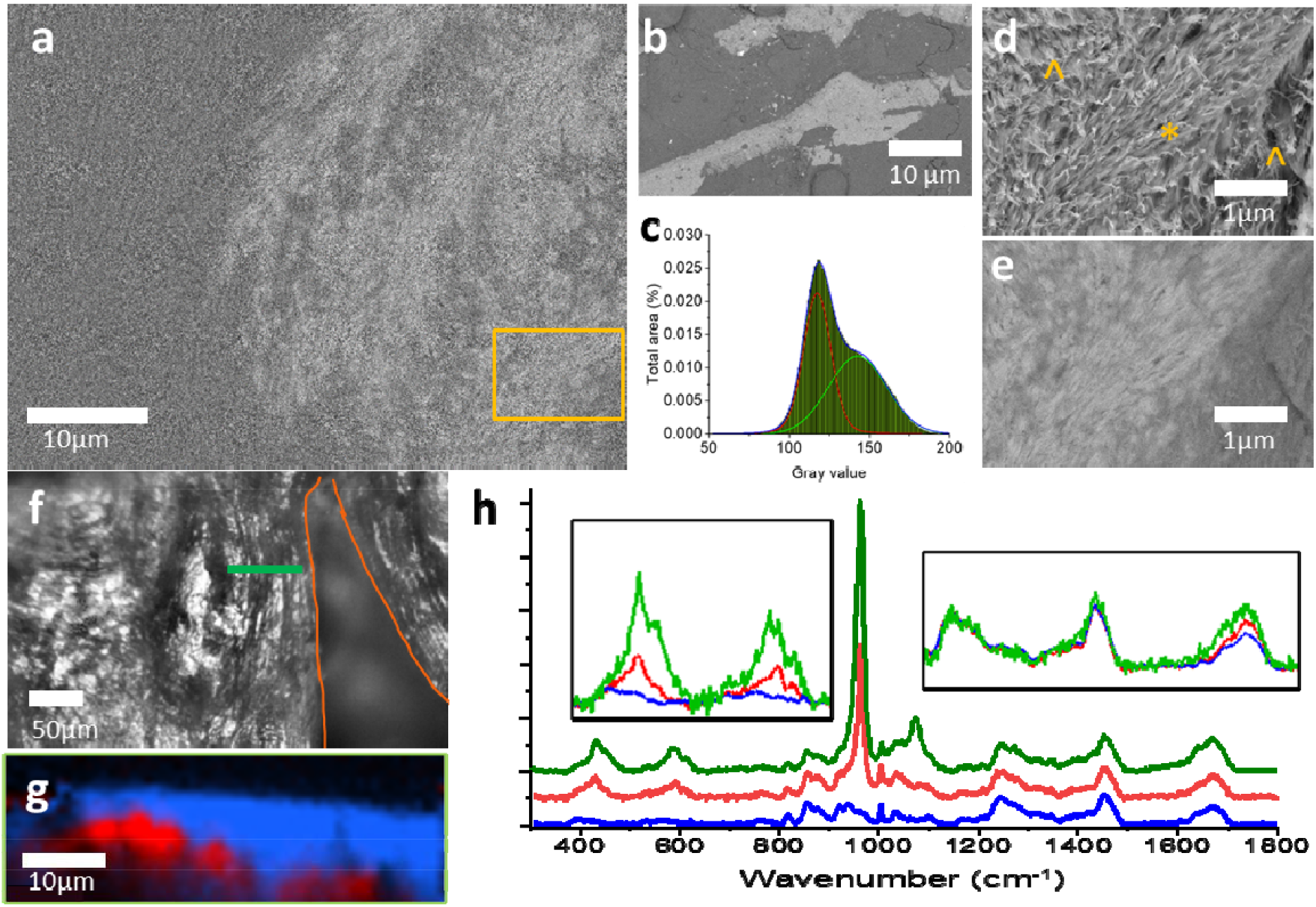
Partially re-mineralized matrix a-b) SEM-BSE image from a) a partly re-mineralized matrix section showing islands of mineralized collagen (light contrast) in an unmineralized matrix similar to what is observed in b) developing mouse bone (see also Figure S7). c) Histogram of the gray values extracted from the BSE image in a) with peak fitting to provide semi-quantitative information on the mineralization progress. Mineralized collagen (green): 52%; unmineralized collagen (red) 48 %. Overlap between peaks shows a wide distribution of mineral densities in the mineralizing matrix. d) High magnification SEM (secondary electron) image showing loose fibrils in an early stage of mineralization both parallel (*) and perpendicular (^) to the imaging plane. e) BSE image of the area in d) showing mineral infiltration follows the fibrils. f) Optical image of the edge of an osteon in the partially re-mineralized matrix. The orange line indicates the border of the osteon with the interstitial bone. g) Raman depth scan (60*µ*m wide; 20*µ*m deep) on the location highlighted in green in (f); g) component analysis identifying two chemically distinct regions. h) Raman spectra extracted from the component analysis in (g). Blue and red spectra correspond to blue and red color maps, respectively. The green spectrum corresponds to the control bone. Spectra are normalized to the orientation independent CH_2_ vibration (1450 cm^-1^). Left inset: Magnification of orientation-independent phosphate vibrations (phosphate ν2 431 and ν_4_ 585 cm^-1^) indicating mineralization initiated in the bottom section (red) of the sample. Comparison with control bone (green) shows the mineral-to-matrix ratio in bottom section is 50% of control bone. Right inset: magnification of the organic matrix vibrations showing only an intensity increase for the orientation-dependent amide I vibration.

On the same sample 3D Raman microscopy was performed using a depth scan on the edge of an osteon (Figure 4f,g), along a line of 60 *µ*m with a scanning depth of 20 *µ*m. Component analysis of the spectroscopic data (see S2) identified both areas of unmineralized (blue) and mineralized (red) collagen (Figure 4g,h). The Raman data confirmed a partial remineralization of the red region (50% compared to control bone) by the analysis of the intensities of the polarization independent phosphate ν2 (431 cm^-1^) and ν4 (590 cm ^-1^) vibrations relative to the amide III (1250 cm^-1^) vibration. A similar phenomenon was reported for fetal mouse bone, where mineral deposition initiated in the bulk of the matrix (Figure 4b, Figure S7 and ref^[9]^). The mineralized collagen domains were present as intergrown 2-5 *µ*m globular structures *within* the matrix, alike the mineral ellipsoids observed in many other studies. ^[8a, 29-30]^

### Mineral development as function of matrix mineral interactions

After establishing that our model reconstructs the mineralized bone matrix in a biomimetic process that requires no activity of osteoblasts or other bone-specific cells, we set out to study the mineral infiltration and maturation as a result of matrix-mineral interactions during bone biomineralization. We used *in situ* 3D Raman microscopy to follow the process in both an osteonal region (Figure 5a,b) and in a region of interstitial bone (Figure 5c,d), i.e. the dense mature bone between the osteons (see Figure 1a).

**Figure 5.**
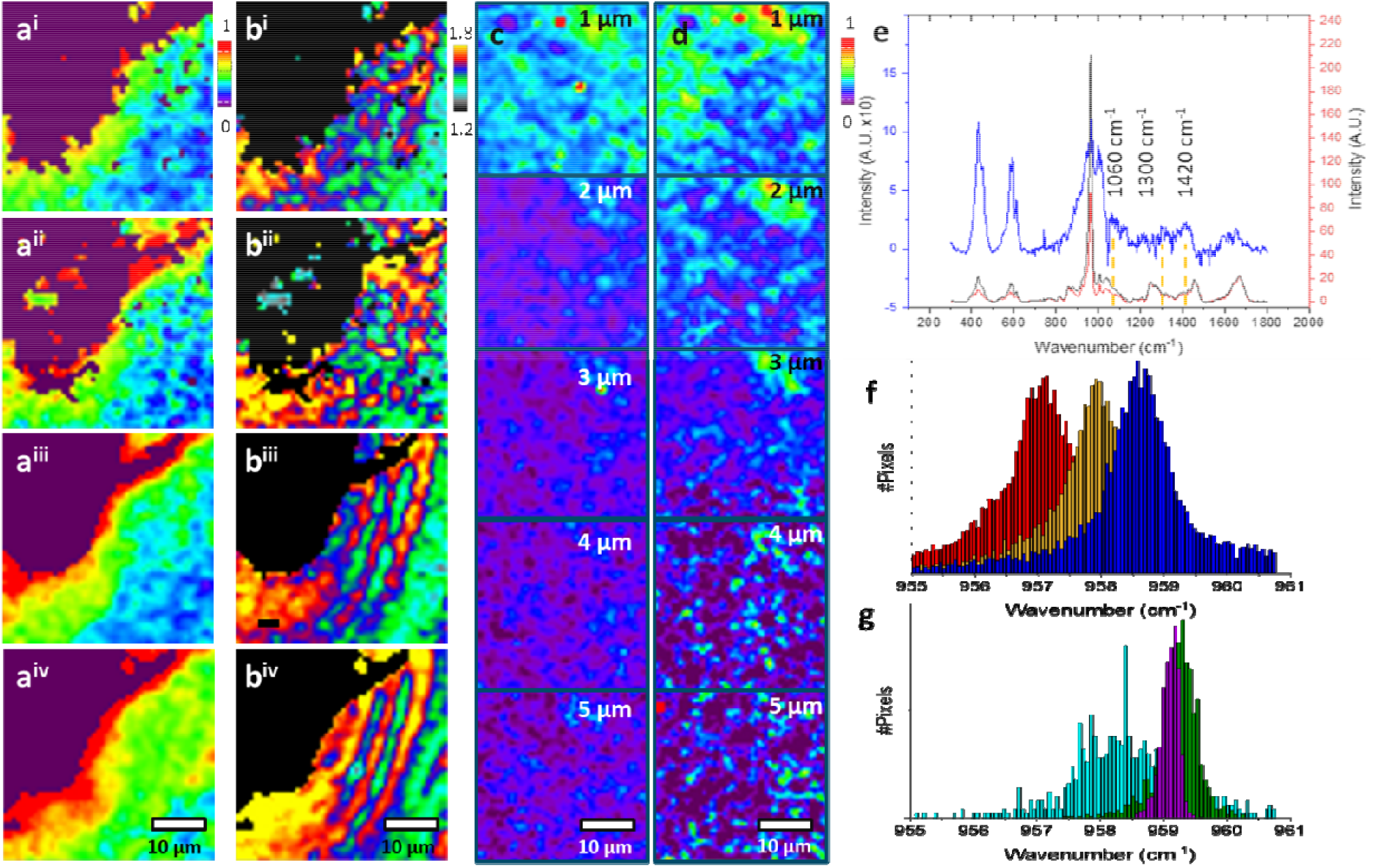
Mineral matrix interactions. a-d) In situ Raman heat maps of re-mineralizing matrix in the (a,b) osteonal and (c,d) interstitial areas’s (see also figure 1a). a,b) Maps were recorded at a depth of 5*µ*m under the sample surface with one day time intervals. a) development of mineral to matrix ratio as determined from the intensity ratios of the (ν2 PO_4_ and CH_2_ regions (431 cm^-1^ and 1450 cm^-1^, respectively) at i) one day, ii) two days, iii) three days and iv) four days of mineralization. The purple area indicates the center of the osteon. b) development of collagen orientation as result of matrix mineral interaction as indicated by the intensity ratio of the orientation-dependent Amide I (1666 cm^-1^) and the orientation independent Amide III (1250 cm^-1^) vibrations at i) one day, ii) two days, iii) three days and iv) four days of mineralization. c-d) Maps recorded at c) day 1 and d) day 9 at different depths below the surface, showing the development of mineral to matrix ratio as determined from the intensity ratios of the (ν2 PO_4_ and CH_2_ regions (431 cm^-1^ and 1450 cm^-1^). e) Average spectra of the pixels showing strongly mineralized collagen, in the interstitial (red) and osteonal (black) regions. Difference spectrum (blue) was obtained after normalization to the Amide III (1250 cm^-1^) peak and correction for the presence of HAp in the phosphate ν1 and ν3 region (900-980 and 1010-1060 cm^-1^, respectively) (See S8). f-g) Histograms of the maxima of the phosphate v1 peaks showing mineral maturation in f) the interstitial region (see also S9), after 24 hrs day (red) and 96 hrs (orange) and 9 days (blue) compared to g) the osteonal region after 4 hrs (cyan) 24 hrs (green), and 96 hrs (purple).

In a region around an osteon, we recorded images at 5 *µ*m below the sample surface, monitoring mineral formation in the bulk of the sample, rather than at the surface of the section. Over the course of 4 days, time resolved Raman maps showed that the mineral infiltration progressed outwards from the Harversian canal (i.e., the center of the osteon) into the organic matrix - in line with the recently proposed vascular access route for mineral transport.^[8a, 31]^ The maps (Figure 5a-d) also showed that natural mineral-to-matrix ratios (Figure 1e,g) were achieved.

In that same period, no mineralization was observed at 5 *µ*m below the surface in the interstitial region (Figure 5c,d lower panel). Investigation of higher layers (1.0 *µ*m, 2.0 *µ*m, 3.0*µ*m below the surface) after 9 days showed that mineralization had progressed slowly, through infiltration from the surface into the bulk, over a depth of only 3*µ*m (Figure 5d), compared to ∼30 *µ*m lateral infiltration in the osteonal region (Figure 5a iv).

Subtraction of the mineralized collagen spectra in the two different bone regions revealed higher polysaccharide signals around 1420 cm^-1^ and 1130 cm^-1^ (Figure 5e and Figure S8c) ^[32]^ in the osteonal region. The concomitant increase at 1660 cm^-1^ (Amide I vibration) indicates these polysaccharides are associated with a protein component and hence suggest higher local concentrations of proteoglycans (PGs) (Figure 5e). The signature peaks in the organic region at 1420 cm^-1^ and 1300 cm^-1^ indicated sulfonation of the carbohydrate chains (see also S4). ^[32]^

Apart from the difference in mineralization rate,^[19a]^ we also noted another significant difference in the mineral development:

If we plot the maxima of the phosphate ν1 vibration for each individual mineral containing pixel in the Raman image, we observe that in the interstitial region after 24 hours most of these maxima center around 957.2 cm^-1^ and over the course of 9 days shift to center around 958.5 cm^-1^, as expected for the frequently reported transformation from a HPO_4_ -based mineral phase (e.g. acidic amorphous calcium phosphate to a purely PO_4_ - based mineral phase (HAp) (Figure 5f, Figure S9a). ^[21b, 33]^ In contrast, the distribution of the phosphate ν1 peak maxima at the osteonal region after 4 hours shows a distribution around 958.2 cm^-1^. Already after 24 hours this distribution had shifted its center to 959.4 cm^-1^ where it remained until 96 hours in the experiment (Figure 5g, Figure S9b). This indicates a much faster transition of the precursor to the final apatitic state in the osteonal region as compared to the interstitial region.

This implies that the maturation of the mineral occurs much faster, beyond the time resolution of the measurement, in the osteonal region as compared to the interstitial region. Given that the same reaction conditions were used for both observations, we attribute the difference in mineral maturation to local differences in the composition and/or structure of the matrix.

For the partially mineralized samples, back scatter electron imaging (Figure 4d,e) had shown that the signal of the mineral follows the organization of the individual collagen fibrils indicating strong guidance of the mineral deposition by the collagen matrix. Raman imaging showed that during the process, the mineral-to-matrix ratio did not exceed the natural levels, demonstrating that the matrix also controls the level of mineral deposition. We note that higher mineral-organic ratios were only observed on the inside of the wall of the Harversian canals in the osteonal regions, where deposits were formed of the pAsp /mineral complex (often referred to as polymer induced liquid precursor-PILP).^[10c, 33-34]^ The pAsp specific peak at 1789 cm^-1^ was not detected in the mineralized matrix. This implies that - in contrast to what is often assumed^[35]^ –the pAsp does not enter the osteoid, but only acts as crystallization control agent that prevents its precipitation outside collagen matrix.^[36]^

Comparison of averaged spectra from the control bone, the unmineralized collagen and the partially mineralized collagen around the osteon (after normalization to the amide III vibration at 1250 cm^-1)^) (Figure 4h right inset), revealed no changes in the matrix related to the subsequent demineralization and remineralization steps, apart from the intensity of the orientation-dependent amide I vibration (1666 cm^-1^) that decreased upon demineralization and increased again during the remineralization process. This indicates changes in the collagen orientation, and imply thereby that mineral-matrix interactions are also a determinant factor in the molecular scale organization of the organic matrix. Indeed, *in situ* mapping of the ratio of the orientation-dependent Amide I and the orientation independent Amide III peaks (Figure 5b) during the infiltration process revealed that the concentric lamellar pattern of the collagen layers in the osteon only re-appeared upon interaction with the mineral. In the setting used here the observed intensity change of the Amide I vibration reflects a change in the organization of the collagen molecules upon interaction with infiltrating mineral.

## 4. Discussion

Above we demonstrate that the cell-free *in vitro* remineralization of demineralized bone matrix provides a physiologically relevant model system to study mineralization processes in the bone extracellular matrix. Demineralization using EDTA was used in order to best preserve the ECM, including matrix associated proteins and GAGs. The model goes beyond other cell free models^[6c, 10a, 10b, 13-14, 35]^ in that it - with high fidelity - reproduces both the structure the chemistry of the bone matrix and that it does this over multiple length scales, i.e. from nanoscale level of the mineralized collagen fibrils, to the sub-millimeter level of the osteons. Moreover, we demonstrate that it also captures intermediate stages of bone biomineralization^[8b, 21b, 37]^ providing confidence that our system faithfully mimics the major pathways of *in vivo* collagen mineralization in bone. Our system further suggests that the presence of intermediate stages and speed of mineral infiltration is in part influenced by the presence of GAGs in the ECM.

Currently the direct dynamic *in situ* monitoring of collagen mineralization in bone has been achieved only using zebrafish and mouse models,^[8a, 21b, 37]^ where ethical considerations, as well as the damaging effect of the applied microscopic techniques to the living tissue are limiting both the duration of the experiments and the maximum obtainable resolution. Other animal models only allow the nanoscale analysis of the mineralizing matrix *ex vivo*, and only provide information on the developmental stage at the timepoint of sacrificing the animal. And although the generation of bone organoids is a exiting new development, they cannot yet provide information on the hierarchical development of the mineralized matrix.^[38]^ In contrast, the current cell-free *in vitro* system allows us to conduct experiments for any duration, and to investigate matrix mineralization *in situ* at multiple length scales. Importantly, in this macroscopic system mineral infiltration is facilitated by the Harversian canals which have been indicated as the natural pathways for the mineral from blood into the tissue.^[8a, 31]^ Beyond what is possible in *in vivo* systems, our system allows us to define a starting point for the experiments, with no ongoing processes at t=0.

The faithful, detailed in vitro reconstruction of the mineralized matrix with the physical composition of bone also verifies our hypothesis that bone matrix mineralization – in contrast to what has been suggested^[7a, 7b]^ - does not require cellular control/involvement after the ECM is fully formed. Rather, it is the composition and structure of the organic extracellular matrix, produced mainly by the osteoblasts, that controls the final organization and chemistry of the material. Moreover, since the composition of the used mineralization solution mimics closely the natural composition of the extracellular fluid, this suggests that indeed direct pathways between the vasculature and the extracellular matrix are enough to provide the tissue with the necessary mineral, providing further evidence for a non-cellular origin of the deposited mineral. ^[8a, 8e, 10a, 10b, 14, 39]^

However, our results do not exclude that osteoblasts produce proteins or other molecules that act as crystallization control agents, similar to pAsp in our system. Indeed for example osteoblasts produce osteopontin^[11]^ and citrate^[40]^ that likely act as crystallization control agents, preventing random crystallization of mineral in the extracellular space.

Nevertheless, it was demonstrated previously that in the absence of pAsp, also fetuin - a protein produced by the liver and the most abundant non-collageneous protein in bone - can fulfill this role^[10a]^. Moreover, we demonstrate that the pAsp does not enter the collagenous matrix with the mineral (see S10). Although these results contrast the results of Jee et al.^[35]^ who used fluorescence experiments to study mineral infiltration in turkey tendon, they are in line with a previous report showing that mineralization of collagen fibrils can also be achieved using neither pAsp, fetuin nor any other crystallization control agent.^[17b]^ Overall our work shows that using an external source of mineral all aspects of the 3D organization of the bone mineral within the collagenous matrix can be recapitulated, and that cellular activity is not strictly needed for this process.

Our model shows a significant difference in the mineralization of the osteonal bone compared to the interstitial bone. Here the interstitial bone, which was low in GAG content, shows a slower progression of mineral maturation and limited mineral infiltration, whereas the osteonal tissues, with higher GAG content showed fast maturation of the HAp crystals and rapid mineral infiltration. Since the regions were exposed to the same mineralization solution, the difference in mineralization behavior must be assigned to variations in the structure and/or composition of the extracellular matrix, where our Raman observations indicate a variation in the GAG composition between the different regions. Although the extracted spectral fingerprint points to a higher abundance of sulfonated GAGs in the osteonal regions, possibly as proteoglycans, contributions of other components to the spectrum, like NCPs to the 1660 cm^-1^ band cannot be excluded.

The interstitial bone shows a lower infiltration of mineral (up to 3*µ*m over 9 days) compared to the osteonal bone (∼30*µ*m over 4 days). We relate our observations to the higher abundance of - at least in part - sulfonated GAGs in osteonal bone. This is in line with recent work identifying higher concentrations chondroitin sulfate (CS) around the osteons compared to the interstitial regions, using enzymatic digestion and histochemical staining.^[19b]^ Recent research has also shown that the most direct interaction between the organic and the inorganic matrix is through glycan interactions,^[18, 41]^ rather than collagen or non-collagenous proteins, which very likely promote mineral infiltration by increasing the wettability of the collagen fibrils.^[42]^ The favorable mineral-matrix interactions most probably also are responsible for the initial confinement of the mineral to the collagen fibrils (Figure 4d,e), as has been observed in several other studies.^[6a]^

Our work also points to a role of the GAGs in promoting the maturation of the mineral. The observed crystalline apatite directly at the mineralization front in the osteonal regions suggests that the mineral associated to the collagen crystallizes much faster in the presence of GAGs than in their absence. Our finding is surprising, as it contrasts several *in vivo* studies that have reported that collagen mineralization with apatite is preceded by the deposition of amorphous or poorly crystalline transient precursor phases.^[21b, 37]^ Here we have to take into account that *in vivo* the mineralizing matrix is the pristine, recently formed osteoid, which is likely different in composition from the mature matrix that is used in the present study. Indeed, several studies have reported the time dependent variation of the GAGs content of the bone extracellular matrix, both prior to the mineralization process^[43]^ and upon aging of the bone matrix. ^[19b]^ Significantly, our work shows for the first time that in bone biomineralization the collagenous matrix not only directs the shape and orientation of the mineral platelets but also promotes their crystallization to crystalline apatite. At the same time, our experiments indicate that the matrix-mineral interactions are not only directive for the mineral formation process, but that the mineral also directs the organization of the collagen. Collagen mineralization has been shown to be a process that involves close packing and dehydration of the collagen molecules, with the developing mineral pushing apart the molecules to make place for the growing apatite platelets.^[6a]^ This imposes contractile pre-stress on the collagen fibrils, thereby significantly contributing to the mechanical properties of the bone ECM.^[44]^ Accordingly, we show that only upon exposure of the collagen to the infiltrating mineral, Raman microscopy reveals the lamellar matrix structure of the osteon. Where the invariant peak shape of the amide I vibration shows that the triple helical structure of the collagen was present throughout the process, the observed increase in this peak implies a re-orientation of the collagen molecules with respect to the incoming laser beam.

Due to their size and the fact that they are cross-linked, we may not expect the orientation of the entire fibrils to change during the mineralization process. Hence, we must attribute this effect to the re-orientation of the triple helical collagen molecules within the fibrils, in line with the expected loss of conformational freedom for the collagen molecules that become locked-in during their embedding within the mineral phase.

## 5. Conclusion

*In vivo* studies toward the roles of PGs and GAGs have been complicated by the fact that PGs fulfill multiple roles in the extracellular matrices (ECM) of different tissues^[20]^ and compensatory mechanisms can counterbalance roles of the deleted PGs and GAGs in knock-out experiments.^[45]^ On the other hand, many *in vitro* systems cannot provide the context of the complex structure of the ECM that significantly determines the activity and action of these biomolecules in different processes.^[13]^

The 3D cell-free *in vitro* model presented here provides a powerful alternative to study matrix-mineralization interactions in bone. It demonstrates that bone biomineralization is both a matrix *driven* and matrix *controlled* physiochemical process, which does not require cellular activity. In particular the combination with in situ 3D Raman microscopy allows the spatial and temporal resolved mapping of the individual but interactive developments of matrix and mineral. Together with electron microscopy observations this confirms earlier observations^[19a]^ that GAGs play an important role in the matrix-mineral interactions which determine how fast, where, and in which form the mineral is deposited, and thereby importantly contribute to the mechanical properties of bone. In this light it will be interesting to explore the value of this model for the investigation of factors that influence the mechanical properties of bone and other mineralized tissues. Indeed, this cell free mineralization approach also holds promise for both mechanistic studies of mineralization related diseases where the mechanically affected matrix also is thought to trigger pathological calcification, such as the rare brittle bone disease osteogenesis imperfecta^[46]^, and heart valve calcification.^[47]^

## Experimental Section/Methods

### Human bone samples

Whole human femurs were retrieved from an anonymous healthy male donor in compliance with European regulations for human cells and tissues (directives 2004/23EC, 2006/17EC and 2006/86EC), and were kindly donated by HCM Medical (Nijmegen, the Netherlands). Femurs were freshly frozen at -20°C, and serologically tested to be free of major pathogens (HIV, hepatitis B/C, HTLV) and microbiological contamination (Clostridium, Streptococcus, fungi). Femur shafts were cut using a hand hacksaw and subsequently trimmed to cubes of approximately 2×2×1 mm on a rotating disc sander under continuous water cooling. Demineralization of bone blocks was carried out in a solution of 9.5% EDTA, in PBS at pH 7.4 at room temperature for 12 days. Each block was submerged in 5ml of demineralization solution. The solution was renewed every workday. From the demineralized bone cubes, slices of 200*µ*m thickness were cut using a Leica VT1000S vibratome using a steel blade for the cutting. Slicing was performed with the speed and vibration frequency set to 3.5 and 3, respectively.

### Collection of ICP-OES samples

Prior to each solution change 1 ml of the solution was collected and stored at -18°C. The remainder of the solutions was acidified with 100 *µ*l of 65% nitric acid and subsequently diluted to 10 ml. After acidification the samples were centrifuged and the solutions filtered to remove precipitation. A reference sample of the demineralization solution was also measured to check for trace amounts of calcium. This was determined to be 0.1589 ppm. For the calculations, this value was deducted from the measured amounts of calcium for the rest of the solutions.

The solutions were analyzed with an ICAP 6000 ICP-OES spectrometer (Thermo Fisher Scientific, Bremen, Germany) equipped with a vertical torch and a radial plasma observation to determine the amount of resorbed calcium from the bone specimens.

### Preparing general remineralization solution

The slices were remineralized using a solution prepared by mixing equal amounts of 6.8 mM CaCl_2_, 4 mM K_2_HPO_4_, 3.4×PBS (containing 465.43 mM NaCl, 9.12 mM KCl, 6.49 mM KH_2_PO_4_ and 29.33 mM Na_2_HPO_4_.H_2_O) and 400 *µg/ml* polyAsp. This resulted in a final concentration of 0.8x PBS solution containing 1.7 mM CaCl_2_, 1.0 mM KH_2_PO_4_ with 100*µ*g/ml polyAsp. All pure materials were dissolved in MiliQ water. Before mixing the pH of the PBS was set to 7.4 for non-carbonated remineralization and 7.1 for remineralization with carbonated solution. For carbonate infiltration the PBS solution was supplemented with 100mM of KHCO_3_.

### Reconstructing intermediate mineralization stages

Sections from the demineralized block obtained as described before were submerged in 1.5 ml of the general remineralization solution. The samples were placed on a rotating stage for a total of 8 days under constant movement at 37 °C. Solution was renewed once after 4 days of incubation. After which the samples were prepared for SEM imaging.

### Preparing solutions for In situ analysis of ECM remineralization

Sections of roughly 200 *µ*m in thickness were cut from the predescribed demineralized slices. 2 sections were stuck to a SEM stub with carbon tape. Samples were exposed to 1x PBS under vacuum for apx. 1 hour to remove air bubbles. The 1x PBS was removed and the stubs were place in a 6-wells plate well. (Biorad). To this well, 12 ml of the earlier described remineralization solution was added at the beginning of the experiment which was renewed daily.

### X-ray imaging

X-ray imaging on the bone samples was performed using a KaVo OP 3D Vision imaging set-up. Images were taken using the Panoramic HD – small settings using kVp 84 at 5 mA. Scanning time was 18.3 seconds.

### Fourier transform infrared spectroscopy

The IR spectra of the bone smaples were generated using a Spectrum One ATR-FTIR (Perkin-Elmer) with a diamond/ZnSe ATR crystal. The spectra were measured in the 400-4000 cm^-1^ wave number range at a resolution of 0.5 cm^-1^.

### Raman spectroscopy

Raman spectroscopy was performed using a WITec alpha300 R confocal Raman microscope (WITec, Ülm, Germany). With excitation laser at 532 nm at a laser power of 40 mW. Imaging was performed at a resolution of 500 nm lateral and 1 *µ*m axial resolution with exposure time of 1 s per pixel. For analysis of the data see S2

### Scanning electron microscopy

SEM samples were first dehydrated using a graded ethanol (EtOH) series with ascending concentrations (v/v) of 50%, 75%, 99% (2x) and ultra-pure EtOH at 4°C, for 15 minutes each on a rocking table. After dehydration the samples were dried with a SPI-DRY critical point dryer.

For scanning electron microscopy (SEM) studies all samples were coated with carbon using a Leica ACE600 sputter coater to a total thickness of 16 nm. Imaging was performed using a Zeiss Crossbeam 550 equipped with a field-emission gun operating at an acceleration voltage of 5kV, a probe current of 67-69 pA and a constant working distance of 10mm.

### Transmission electron microscopy

Following demineralization, the bone blocks were infiltrated overnight with sucrose solution for cryoprotection. subsequently, they were positioned on a mounting pin and plunged into liquid nitrogen. The frozen specimens were then sectioned into 210 nm thin sections with a cryo-microtome, after which the sucrose was washed out using PBS. Finally, the samples were transferred to Quantifoil® TEM grids (EMS Diasum, PA, USA).

Samples were remineralized on grid using by using 50 *µ*l of remineralization solution for 2 hours. After this time the solution was blotted and the samples were washed by placing the grids on droplets of miliQ water twice, to remove excess salts. After which the samples were dried under vacuum. TEM imaging was performed using a A JEOL JEM-2100 system (Jeol Ltd., Tokyo, Japan) equipped with a LaB6 filament was used for imaging at 200 kV. Images were recorded with a Gatan 833 Orius camera (Pleasanton, CA, USA).

## Acknowledgements

We thank Adi Ben Shoham (Weizmann Institute of Science) for allowing us to use images of foetal murine bone development to be compared to our in vitro bone model. We also thank Neta Varsano (Weizmann Institute of Science^[48]^ for generating the bone and collagen illustrations. HCM medical (Nijmegen, the Netherlands) for providing the human bone sample. We acknowledge the contributions of Michiel Beurskens in developing the sample demineralization. The project was supported by an European Research Council (ERC) Advanced Investigator grant (H2020-ERC-2017-ADV-788982-COLMIN) to N.S. A.A. was in part supported by a VENI grant from the Netherlands Scientific Organization NWO (VI.Veni.192.094).

