## Supporting information for "A 3D cell-free bone model shows collagen mineralization is driven and controlled by the matrix"

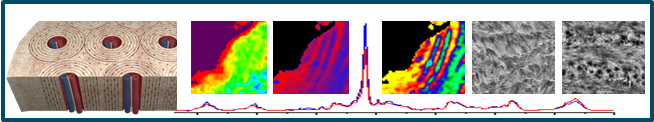

Through in vitro remineralization of demineralized bone matrix we restore all hierarchical levels of mineralization from the apatite nanocrystals to the structure of the osteon, without the need of cellular activity. We show that matrix-mineral interactions and the presence of glycosaminoglycans play important roles in directing and controlling mineral development.

Supporting Information for

A 3D CELL-FREE BONE MODEL SHOWS COLLAGEN MINERALIZATION IS DRIVEN AND CONTROLLED BY THE MATRIX

Robin H.M. van der Meijden,^1,2^ Deniz Daviran^1,2^ Luco Rutten^1,2^, X. Frank Walboomers^3^, Elena Macías-Sánchez^1,2,4^, Nico Sommerdijk^1,2 *^ Anat Akiva^1,2*^

**S1: Progression of bone demineralization**

Demineralization of bone ECM was analyzed using inductively coupled plasma optical emission spectroscopy (ICP-OES), X-Ray imaging, IR-spectroscopy and scanning electron microscopy (SEM) imaging. ICP-OES was performed as described in the materials and methods sections. In short, during the demineralization the extraction solution was renewed daily. Small aliquots of the removed solution were used for ICP-OES measurements to determine the amount of extracted calcium. Figure S1a, shows that over a period of 12 days 99.7% of all available calcium ions were removed from the matrix.

X-ray imaging shows that the mineral is indeed removed through the entire bone sample after 14 days, by the loss of contrast in the image of the demineralized sample compared to the control sample (Figure S1b). Additionally, attenuated total reflection – infrared spectroscopy (ATR-FTIR, Figure S1c) analysis shows that after demineralization (black ), the phosphate ν_3_ vibration visible in the bone spectrum (blue) at 1020 cm^-1^ is lost. The demineralized bone spectrum now resembles a pure collagen sample (red spectrum) rather than bone sample. Finally, SEM imaging (Figure S1d) shows that the demineralized matrix consists of loose collagen fibrils. The absence of back scatted signal (Figure S1e) indicates complete removal of the mineral even at the level of the single collagen fibril.

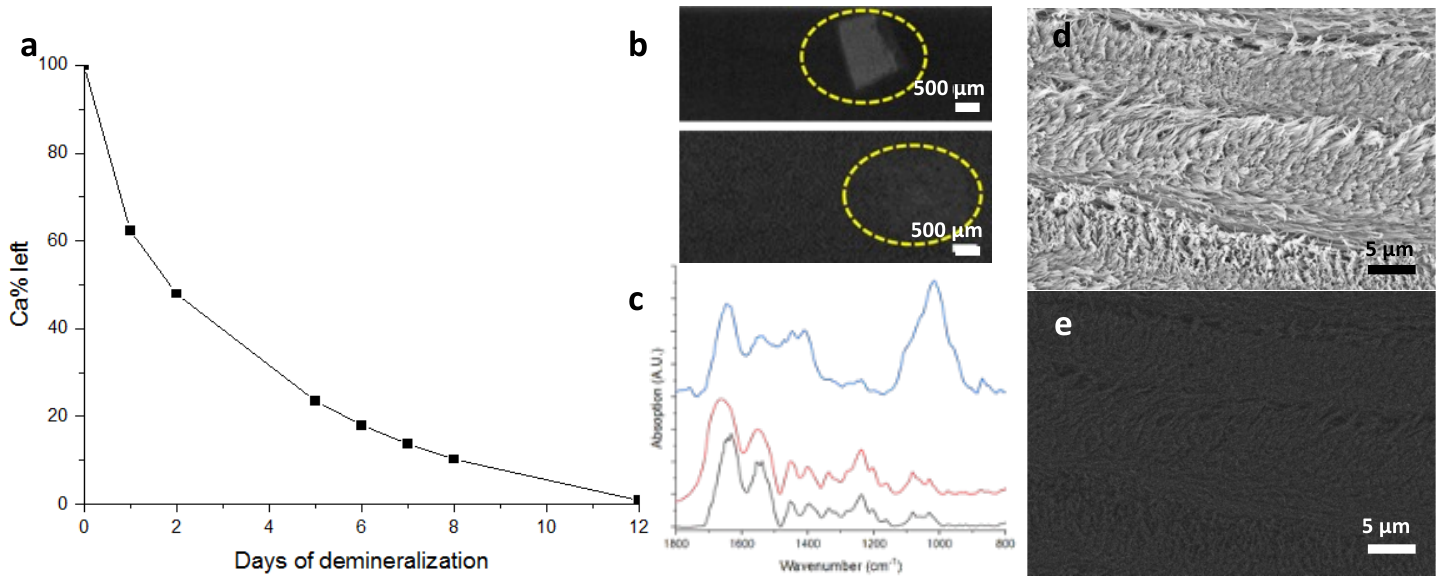

Figure S1 Development of matrix demineralization. a) ICP-OES results showing the gradual decrease of mineral in the bone tissue during EDTA demineralization over a period of 12 days. At the end of day 12, 99.7% of calcium is removed. b) X-ray images of the decalcified matrix before (top) and after (bottom) 14 days of demineralization. The mineralized bone piece shows a gray contrast, while the demineralized bone shows no contrast compare to the background. The sample is marked with dashed yellow ellipsoid. c) ATR-FTIR spectra of bone (blue), collagen (red) and demineralized matrix after 14 days (black). The bone spectrum shows a clear peak at 1020 cm^-1^ indicating the presence of apatite that is missing in both the demineralized and pure collagen spectra. Spectra were recorded on thin sections of human bone, bovine collagen and demineralized human bone, respectively. d,e) SEM images of a demineralized collagen section. d) Secondary electron microscopy (SE) image showing loose collagen fibrils and e) BSE image showing low contrast, both indicating removal of the mineral phase.

**S2:** **Analysis of Raman data**

***General***

Spectroscopic images were all cropped to a size of 50x50 µm and manually aligned to account for drift between different measurements. The spectroscopic data were truncated between 300 and 1800 cm^-1^ and background corrected using the shape function (filter size 300) provided in the Project 5 software (WITec). Data on specific peaks were obtained using the Project 5 software. For peak ratios the area under the curve was extracted for each peak in each pixel according to the parameters in table S1. The values of the peak ratios were determined from these values.

*Table S1: Parameters used for determining area under the curves for all specific features in a bone spectrum.*

| Peak | Position (cm^-1^) | Width (cm^-1^) |
| --- | --- | --- |
| Phosphate ν2 | 431 | 80 |
| Phosphate ν4 | 590 | 80 |
| Phosphate ν1 | 960 | 40 |
| Amide III | 1250 | 80 |
| ~CH_2_ | 1450 | 80 |
| Amide I | 1666 | 120 |

The value for the peak center of the phosphate ν1 peak was determined by using the center of mass value for this peak, taken at the position 960 cm^-1^ over a 40 cm^-1^ region. Histograms were created from these data by plotting the values of the position from 957 cm^-1^ to 963 cm^-1^ divided over 100 bins of equal width. To account for different levels of mineralization during mineral development, a pure collagen spectrum was subtracted to minimized contributions of the proline vibration at 940 cm^-1^ after normalizing both spectra to the amide III (1250 cm^-1^) peak. Removal of non-collagenous contributions to the chemical images presented in this work was done by creating a mask of proline (858 cm^-1^) containing spectra. This excludes the pixels which contain spectra of pure water or poly-aspartic acid, while maintaining the pixels that show spectra of (un-)mineralized collagen.

***Component analysis***

Component analysis was performed to identify the different regions of interest in the sample. Components were identified based on their spectral markers and divided into 4 groups: water, poly-aspartic acid, unmineralized collagen and mineralized collagen. To obtain the quantitative information of the mineralized collagen in each section, a mask was created of the mineralized collagen component by selecting the pixels which have a component value of >100 for this component.

To obtain rotation dependent spectra of mineralized collagen, 12 single point spectra of a control sample were obtained on the same location, while rotating the orientation of the laser, using 10 mW excitation laser (532 nm), 5 s exposure time and 6 accumulations per spectrum. The halfwave plate of the spectrometer was rotated 15 degrees resulting in a 30 degrees shift of the laser rotation per spectrum. Spectra were all normalized to the Amide III vibration and plotted in Origin (Origin Pro 2019b). Where a 3D visualization could be made (Figure S6b,c).

To create a profile plot of the orientation of the re-mineralized matrix (Figure 3e), the processed dataset was normalized to the Amide III (1250 cm^-1^) intensity. From this line sections were taken of 15 µm in length, perpendicular to the osteon direction, along which the area of both the mineral and collagen peaks were determined. The values per pixel along this line were transported to the Origin data analysis software (OriginPro 2019b) and plotted in a 2-axis plot to allow for an overlay of the spectra. The periodicity and width of the lamella was determined by fitting a cosine function to the extracted data using the Origin 2019b software and taking half the period of the fitted cosine to be taken as the width of a single lamella.

***Extracting the GAG-spectrum***

To extract the GAG spectrum from the obtained in vivo data, first an average spectrum of the mineralized matrix was generated by creating a mask of the mineralized collagen based on component analysis of the full dataset. Subsequently an average spectrum of the selected area was produced using the Witec project 5 software, for both the interstitial (Figure 5f, red) as well as the osteonal (Figure 5f, black) areas. These spectra were normalized to the Amide III mode and subsequently the interstitial average was removed from the osteonal average (Figure 5f blue + Figure S8c).

Since the average level of mineralization was higher in the osteonal section compared to the interstitial section, these contributions had to be corrected for. This was done by first creating a deconvolution of a pure hydroxyapatite spectrum (rruff-database, [https://rruff.info/hydroxylapatite/R060180](mailto:)) to identify the positions of the phosphate ν _1_ and ν_3_ sub bands. Subsequently, bands were fitted to the subtracted spectrum, having the same positions as determined for hydroxyapatite, and maintaining the same relative intensities and peak width when comparing phosphate ν_1_ and ν_3_ contributions. These fitted peaks were removed from the subtracted spectrum, to result in the corrected spectrum (Figure 5e, blue) that was used for identification of the compositional difference in the matrix in the different regions.

**S3: Mineral-to-matrix ratio before/after remineralization**

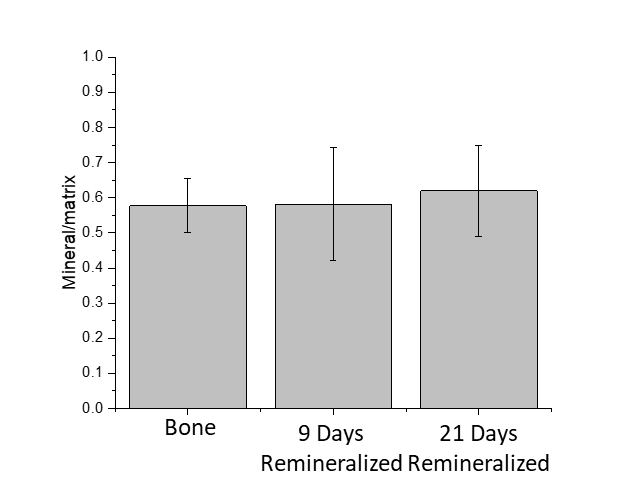

**Figure S3** Average mineralization level and standard deviation (n=6400) of comparable areas from control (bone) and remineralized matrix samples. The mineral-to-matrix ratio is calculated based on the peak areas of the PO_4_ _ν2_ (431 cm^-1^ ) and CH_2_ vibrations (1450 cm^-1^ ).

**S4: Overlap in Raman spectra of GAGs with other bone matrix components**

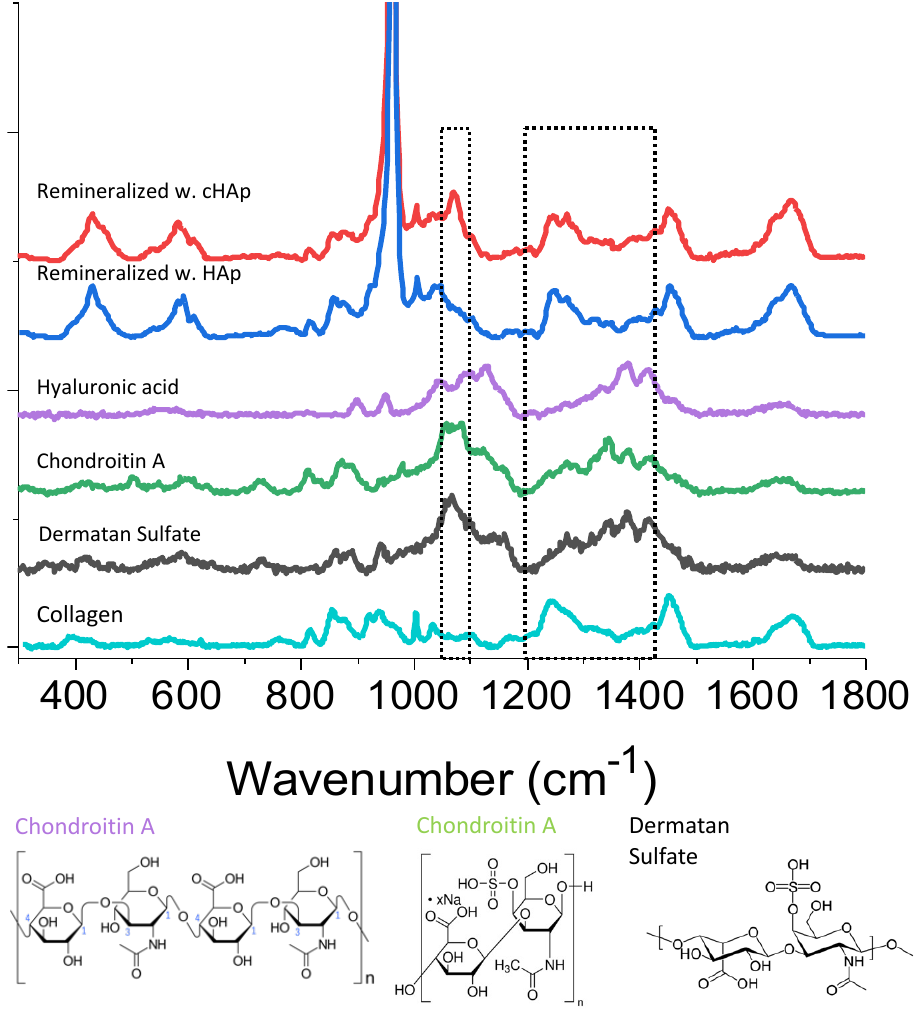

***Figure S4*** *Comparison of Raman spectra of different bone matrix components showing the overlap of the carbonate signal (1030-1090 cm^-1^) from collagen mineralized with cHAP (red) with the signatory peaks of GAG glycosylic bonds around 1060 cm^-1^ as well as the overlap of the collagen amide III peak (1255 cm^-1^) with the characteristic GAG signals in the 1200-1300 cm^-1^ region, as exemplified for hyaluronic acid (purple), chondroitin A (green), dermatan sulfate (black).*

S5: Overlay of optical and SEM micrographs with Raman imaging orientation data

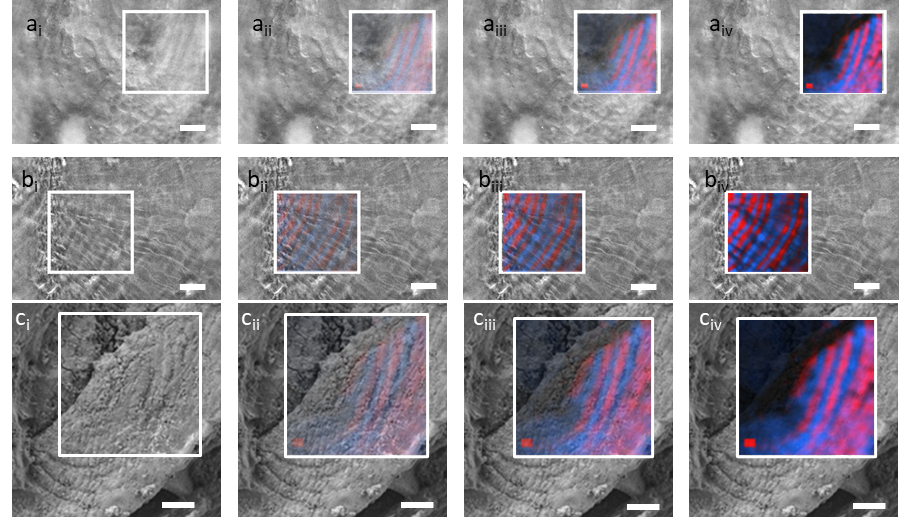

**Figure S5** a-c) Overlay of the Raman microscopy images with optical and scanning electron micrographs as in figure 3h-j with decreasing transparencies for the Raman micrographs (i-iv). a) Overlay of Raman and optical micrograph of bone, b-c) Overlay of Raman with b) optical micrograph and c) SEM image of the remineralized matrix. Scale bars 10 µm

**S6: Orientation dependence of the mineral/phosphate ν_1_ and collagen Amide I Raman spectral signatures.**

The orientation dependence of the phosphate ν_1_ (959 cm^-1^) and the Amide I (1660 cm^-1^) was analyzed using polarized Raman spectroscopy.^[26]^ When the laser is rotated by 90° relative to the collagen orientation, a clear change in both spectra is observed. The phosphate ν_1_ peak increases in intensity when the orientation goes from parallel to perpendicular relative to the fibril, whereas the Amide I peak decreases following the same change (Figure S6a). In order to understand whether this change is also continuous over smaller intervals, an experiment was performed where the laser was rotated with 30° intervals over a total of 180°. Before plotting the spectra were all normalized to the polarization independent Amide III (1250 cm^-1^) peak. This resulted in a continuous change of both the phosphate (Figure S6b) and collagen (Figure S6c) peaks, which followed the same reciprocal relation as shown for the extreme case in Figure S6a. This demonstrates that the shift is continuous and can be used to follow the orientation of the mineralized fibrils.

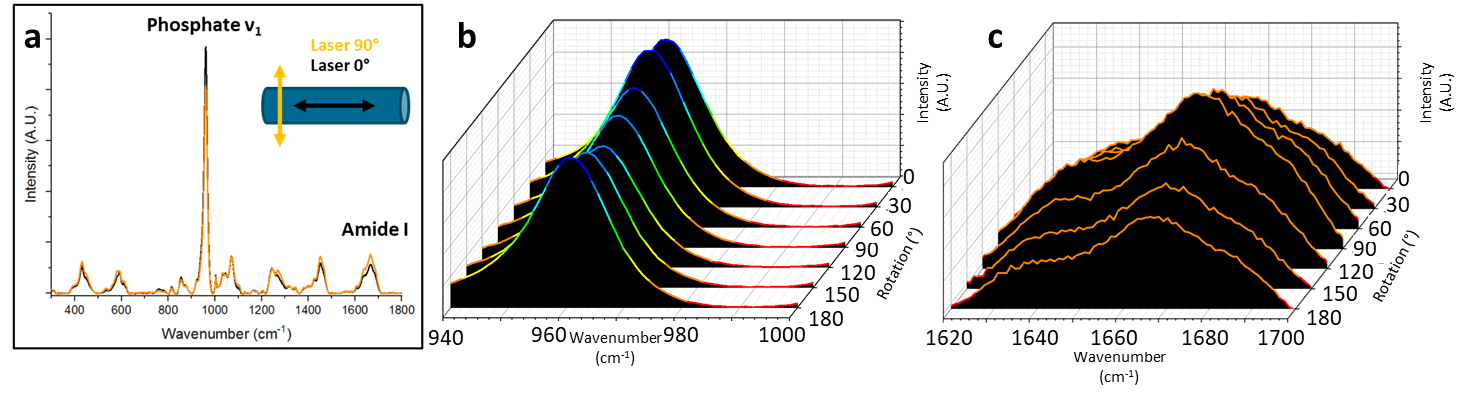

**Figure S6** Orientation dependence of the Raman signal of bone a) spectra of bone tissue with the laser oriented parallel (black line) and perpendicular (yellow line) to the collagen fibril direction, showing the dependence of the Phosphate ν_1_ and Amide I vibrations on the relative orientation, as well as their reciprocal dependence. b) Change in the phosphate v1 peak intensity normalized to the Amide III vibration, as a function of the rotation of the laser. When the angle between the laser and the collagen changes, the signal shows a continuous change as well, being higher in parallel orientation (0°, 180°) and lowest in perpendicular orientation (90°). c) Change in the Amide I peak intensity normalized to the Amide III vibration, as a function of the rotation of the laser. When the angle between the laser and the collagen changes, the signal shows a continuous change as well, being lower in parallel orientation (0°, 180°) and higher in perpendicular orientation (90°).

S7: In vitro mineralization mimics foetal bone mineralization.

To understand how our mineralization process compares to the mineralization process in vivo, we studied the intermediate stages of mineralization by exposing the demineralized matrix to a solution deficient in mineral precursor, to limit the development of the mineralization.^[28]^ These samples were subsequently imaged using SEM microscopy using the backscattered electron detector and compared to images from developing murine bone (courtesy Adi Ben Shoham, Weizmann Institute of Science, experiment was cunducted as part of the study presented in reference^[8c]^). In both samples we observe a patch-like mineralization pattern throughout the sample, with dense mineralization (ligh contrast) in the bulk of the material. These results indicate that our system is capable of recreating the intermediate stages of mineral infiltration.

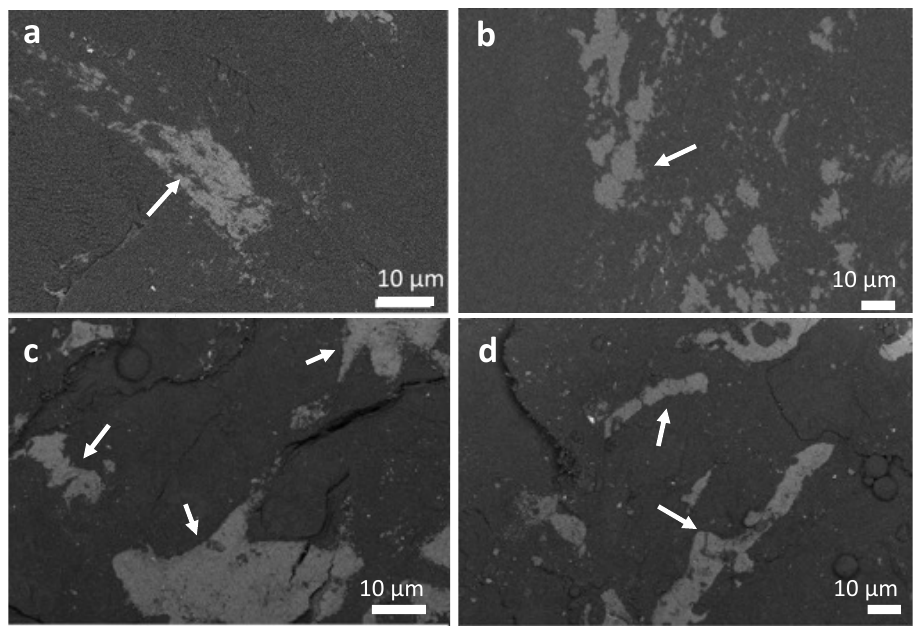

**Figure S7** Backscattered images of a remineralized bone matrix and murine bone mineralization model (E-16). a,b) backscattered electron images from different locations after 8 days of remineralization in a precursor limited solution. c,d) cryo SEM freeze fracture backscattered electron images of forming bone tissue in the epiphyseal growth plate at the proximal tibia of a foetal mouse at embryonic stage day 16. Arrows are pointing at mineral.

S8: Preparing Raman difference spectra for comparative analysis

To allow for comparison of the spectra after subtraction of the remineralized osteonal and interstitial bone regions (Figure 5), we removed the contributions of the mineral from the subtracted spectrum, since the osteonal bone reached a higher level of average mineralization compared to the interstitial bone.

For this we first performed a deconvolution of the phosphate v_1_ and v_3_ peaks in a reference hydroxyapatite spectrum from the rruff database (Figure S8a). This gave us the positions and relative intensities of the two vibrational modes, that we then use to fit to the subtracted spectrum. The peaks identified in the pure hydroxyapatite spectrum were subsequently fitted to the subtracted spectrum (Figure S8b), keeping the positions the identical and keeping the relative areas and peak widths the same as in the fitted spectrum. These peaks were subsequently subtracted from the original subtracted spectrum to result in the corrected spectrum (Figure S8c). Here we see a clear removal of both the vibrational modes of phosphate, leaving only the contributions of the organic matrix.

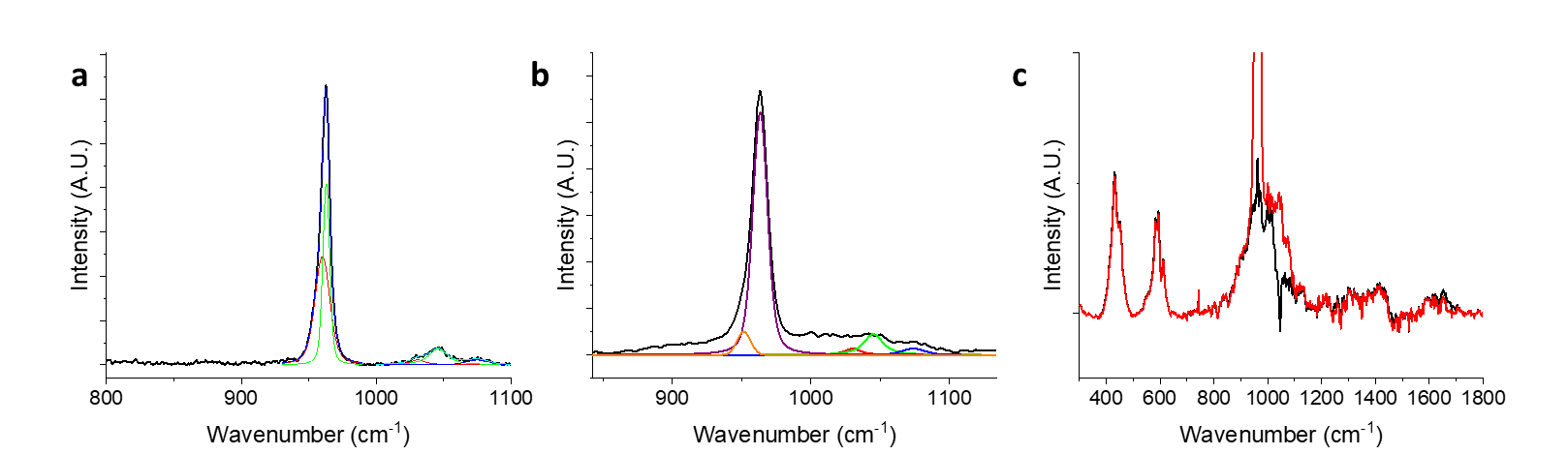

**Figure S8** Deconvolution of the subtracted spectrum. a) Fitting of the sub-bands of the phosphate ν_1_ and ν_3_ vibrations of a hydroxyapatite reference (rruff). This fitting resulted in identification of 5 peaks at 960, 962, 1030, 1045 and 1074 cm^-1^. b) Fitting of the phosphate sub-bands (colored peaks) at the same positions in the subtracted spectrum keeping relative intensities the same as identified in (a). c) Comparison of the subtracted spectrum (red) and the corrected spectrum (black) after removal of the phosphate bands obtained in (b).

**S9: In situ analysis of mineral maturation during matrix mineralization.**

To understand the relation between mineral infiltration and mineral maturation, the central positions of the phosphate ν_1_ peak were plotted as maps for both the interstitial (Figure S9a) and osteonal (Figure S9b) areas. For reliable analysis the position of the phenylalanine was set to be at 1003 cm^-1^. Before the data was plotted a pure collagen spectrum was subtracted from the full dataset, to avoid possible interference of the collagen proline signature at 940 cm^-1^ on the phosphate signal at different levels of mineralization. Plotting the maps of the infiltration process for both samples shows that in the interstitial area, the crystallinity of the mineral develops gradually, starting at 957.2 cm^-1^ after 1 day of infiltration and slowly progressing toward 958.5 cm^-1^ over a period of 9 days (Figure S9a). The same procedure shows that in the osteonal region already during the first day, the crystallinity of the mineral reached its mature form as indicated by the position of the phosphate ν_1_ peak at 959.2 cm^-1^. Only at the mineralization front we observed lower levels of crystallinity which progressed towards a fully crystalline hydroxyapatite phase in only 4 days (Figure S9b).

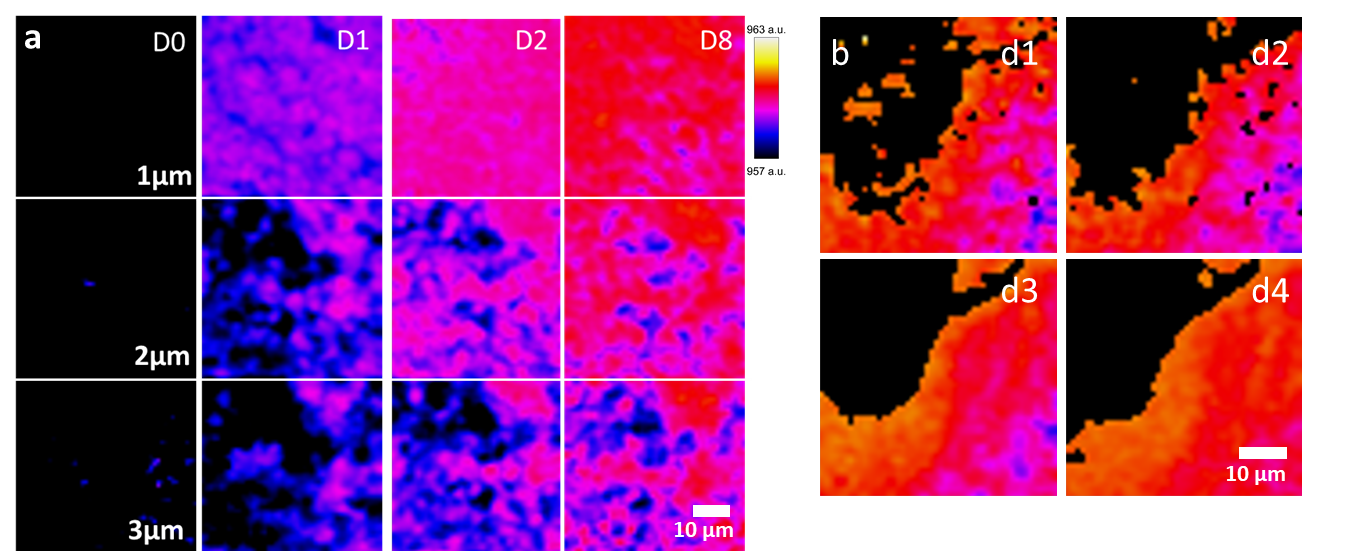

**Figure S9** Visualization of the phosphate ν_1_ peak shift during mineralization infiltration. a) development of the mineral crystallinity as a function of time (horizontal) and depth (vertical) during the in vivo mineralization experiment, in the interstitial bone. We observe a shift from a dark purple to a red color over the period of 8 days, during the mineralization process, indicating a peak shift from 957.2 to 958.5 cm^-1^. b) Development of the mineral crystallinity during a four day period in the osteonal bone. Here we observe that the mineral immediately shows mature hydroxyapatite signature with the phosphate peak at 959.2 cm^-1^ in the mineralized area, only showing a light pink border, indicating less mature mineral, at the mineralization front.

**S10: Lack of infiltration of pAsp in the collagen matrix**

To validate that the pAsp does not infiltrate the matrix, we have performed depths scans on the surface of a washed sample which has been remineralized for 22 days. In this sample we clearly see infiltration of mineral in the collagen, together with a PILP precipitation on the surface of the sample. These results show clearly the detectable signal of the poly-aspartic acid at 1789 cm^-1^ in the PILP precipitation, but not in the collagen matrix, when normalized for the mineral signal. This shows that indeed the pAsp does not infiltrate the matrix as the mineralization proceeds.

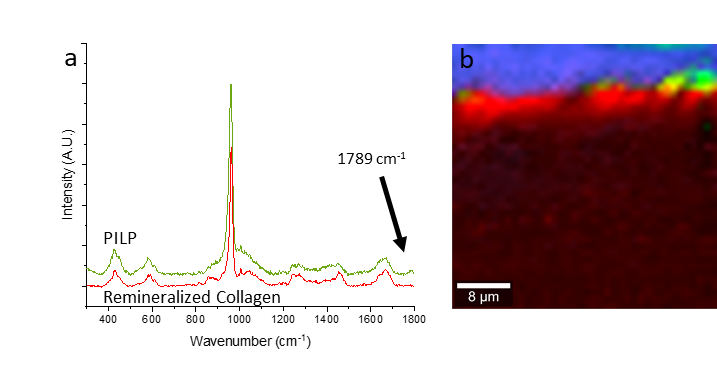

**Figure S10** Raman a) spectra and b) map of the PILP layer deposited on the inner surface of the osteonal matrix (green) and of the mineral deposited inside the collageneous matrix (red). Arrow in a) indicates the pAsp COO vibration at 1789 cm^-1^ that is the main difference between the two spectra. The map shows there is no pAsp inside the remineralized matrix, only on the inside surface of the osteon.
